# Immunophenotyping of HER4/YAP axis in breast cancer brain metastasis

**DOI:** 10.1101/2020.01.29.924118

**Authors:** Priyakshi Kalita-de Croft, Malcolm Lim, Xavier M de Luca, Bryan W Day, Jodi M Saunus, Sunil R Lakhani

**Affiliations:** Faculty of Medicine, The University of Queensland, Brisbane, QLD 4029, Australia; Pathology Queensland, The Royal Brisbane and Women’s Hospital, Brisbane, QLD 4029, Australia; Dept of Cell and Molecular Biology, Sid Faithfull Brain Cancer Laboratory, QIMR Berghofer Medical Research Institute, Brisbane, QLD 4006, Australia; Faculty of Health, Queensland University of Technology, Brisbane 4059, Australia

**Keywords:** Breast cancer, brain metastasis, HER4, clinicopathological features, biomarkers

## Abstract

Brain metastasis (BrM) is a devastating diagnosis for patients with breast cancer. Identification of immunophenotypical features specific to BrM is crucial for developing effective new therapeutic interventions. Yes-associated protein (YAP) mediates downstream effects of the neuregulin receptor, HER4, on breast cancer cell proliferation *in vitro*. HER4 is frequently over-expressed in BrM, but the relationship with YAP has not been investigated. This study examined the HER4/YAP axis in patient samples (n=41) from matched primary breast and metastatic brain tumours. Immunohistochemistry analysis revealed HER4 was highly phosphorylated in BrM compared to matched primary tumours (p=0.0022), and this was strongly associated with expression and phosphorylation of dimerization partners HER2 and HER3 (p<0.0001). When compared to a general breast cancer cohort (n=373), we found HER4 activation to be brain metastasis specific (p<0.0001). However, YAP was frequently phosphorylated in both HER4-activated primary breast tumours and brain metastases, suggesting that pro-proliferative YAP signaling is not a major consequence of HER4 activation in these tumours. These results display the complexity of expression of the HER family receptors and the downstream pathways in BrM and suggest simultaneous targeting of multiple receptors might be more advantageous.

## Introduction

Relapse of breast cancer (BC) in the brain leads to significant morbidity and a poor life expectancy, generally less than two years from diagnosis^1^. The exact incidence of brain metastases (BrM) is difficult to determine because it is not routinely documented in patients with disseminated disease, nor screened for in asymptomatic patients, but the most recent epidemiological data indicate high incidence of up to 40% of intracranial metastases in cancer patients^1–4^. Moreover, with prolonged overall survival of cancer patients due to good systemic control^5^, the incidence of BrM is increasing^6^. Treatment modalities can include stereotactic radiosurgery, surgical resection, focused external beam radiosurgery, whole-brain radiotherapy and conventional chemotherapy^7^. These interventions can improve quality-of-life and overall life expectancy, but are rarely curative. There is a dearth of molecularly-targeted options for systemic therapy. Understanding the vulnerabilities of BrM is crucial to developing new targeted therapies. One way to approach this is to identify molecular features that set BrM apart from their parent primary cancers.

We and others have previously shown that members of the human epidermal growth factor receptor (HER) family of receptor tyrosine kinases are pivotal in the pathogenesis of BrM and treatment resistance, with HER2 and HER3, the most common oncogenic partners, playing a role in colonization of brain metastasis^8–10^. Mechanistically, these receptors induce pro-tumorigenic effects through ligand-dependent activation. Neuregulin ligands (known to be abundant in the brain)^11^ bind to the HER3 extracellular region, resulting in homo or hetero-dimerization with HER2 and consequent recruitment of downstream signaling proteins that bring about changes in transcription, growth, survival and proliferation^12, 13^. HER4 is also receptive to a broad range of neuregulin ligands, can form heterodimers with other family members^14^, and is frequently over-expressed and activated in BrM^9^, suggesting it may be an important player in BrM development.

HER4 has nine known isoforms, and is unique amongst HER family members through its ability to undergo juxta- and intra-membrane proteolysis, releasing an intracellular domain (ICD) with independent transcriptional modulating functions^15–21^. The ICD of HER4 has been shown to induce yes-associated protein (YAP)-regulated genes^22^, and directly interacts with YAP in nuclei of breast cancer cell lines^23^. YAP is a regulator of multiple cellular functions including proliferation, differentiation and survival, with a growing body of evidence pointing towards a role in physical microenvironment recognition^24^. The canonical mechanism of YAP function involves Hippo signaling, which is engaged after phosphorylation and nuclear translocation of YAP. In mammals, this is opposed by LATS1/2, which phosphorylates YAP resulting in cytosolic retention, degradation and reduced expression of downstream targets. Hence nuclear-cytoplasmic shuttling of YAP is central to the control of its transcriptional activity and currently, it is known to be active when not phosphorylated.

Based on existing links between HER4 and BrM^25, 26^, and HER4 with YAP signaling^27–29^, we postulated that frequent over-expression and activation of HER4 in BrM from breast cancer could be associated with YAP activation. Since inhibitors of both HER4 and YAP are currently being assessed in clinical trials^30–36^, and there is an urgent need for new therapeutic options in BrM, the aim of this study was to analyze expression and activation of HER4 and YAP in human BrM. Our overall goals were to investigate this previously unexplored axis as a relevant therapeutic target.

## Materials and Methods

### Tissue microarrays (TMA) and histopathology

We used the Brisbane Breast Bank breast to brain metastasis (BBM) cohort, a resource comprising of formalin-fixed paraffin-embedded breast/brain tumours from patients undergoing surgical resection at the Royal Brisbane Women’s hospital (RBWH). Ethical approval from the Human research ethics committees of the RBWH and University of Queensland was obtained prior to commencement of the study (2005000785/HREC/2005/022). We had 26 matched pairs and in total there were 41 breast cancer samples and 50 brain metastases samples, 5 cases had recurrent brain metastases and we had no access to the primary tissues for two cases. The histopathological review of all the cases was conducted by an experienced pathologist (S.R.L); we selected the commonly used diagnostic breast tumour biomarkers implicated in the prognosis of breast cancer (i) Hormone receptor (HR) and (ii) Human Epidermal Receptor 2 (HER2) and we employed the same scoring criteria for the prognostic biomarkers as previously described^37^. Majority of the primary breast cancer cases were triple-negative (TN) (42%), followed by HER2+ (34%) and rest were HR+ (23%). Brain metastasis samples followed a similar trend with the triple-negative subtype comprising of 43% followed by HER2+ (35%) and HR+ (20%). Tumours were sampled on TMAs for biomarker studies and we undertook morphological and immunohistochemical assessment of matched pair breast to brain metastasis samples. Hematoxylin and eosin staining confirmed the presence of tumour tissue within the TMA cores (Suppl fig 1A). We also made a comparison of our proteins of interest in a general breast cancer cohort, the Queensland follow-up cohort (QFU), collected from patients undergoing resection at the RBWH between 1987 and 1994. This cohort was subdivided into the diagnostic biomarker criteria (Oestrogen receptor -ER+ve, HER2+ve and triple-negative disease). It comprises of 373 cases of primary breast cancers; 75% cases Oestrogen receptor (ER+), 10% cases have ERBB2 amplification and is described in detail elsewhere^37^. A list of antibodies and the conditions used are provided in table 1.

**Table 1.**
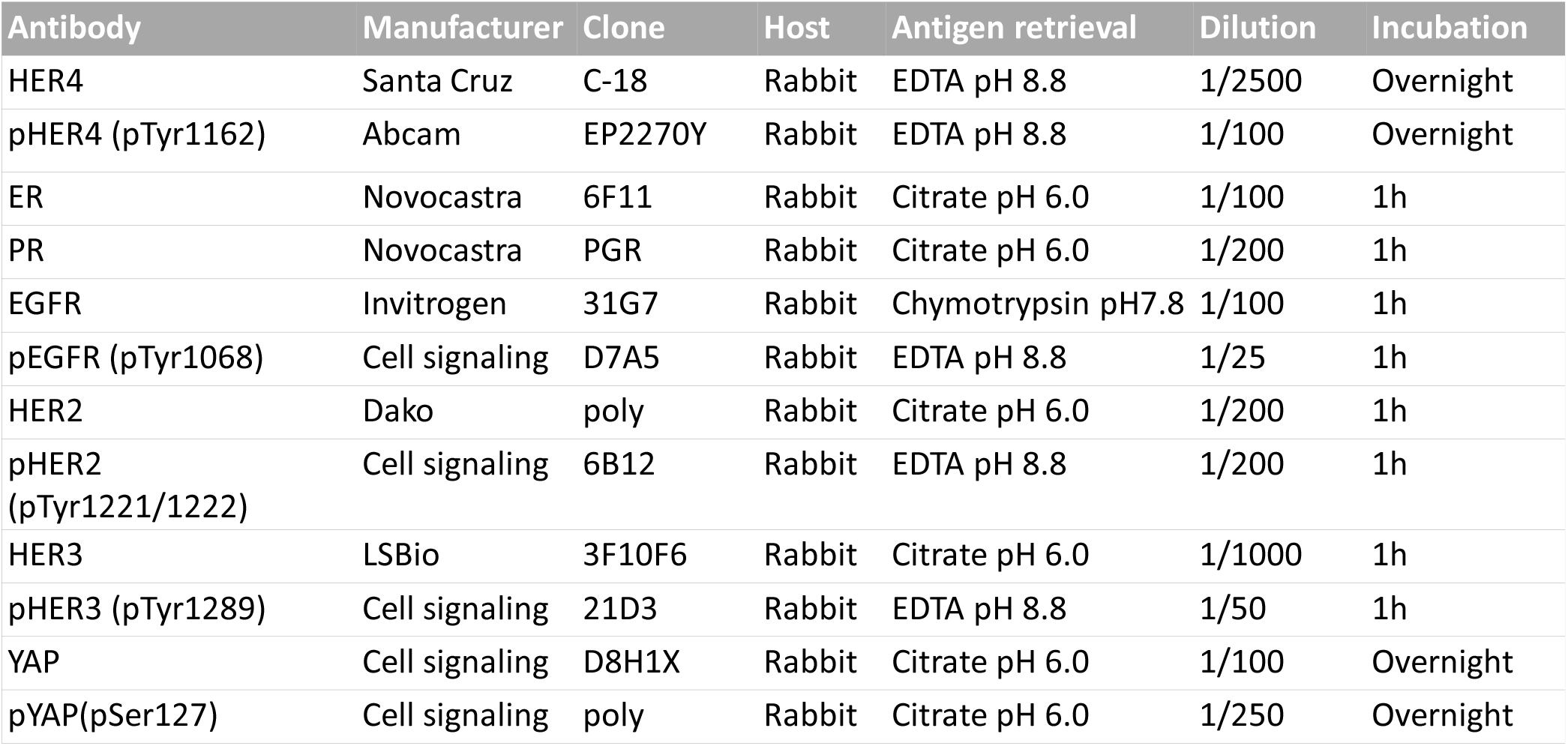
Antibody details and incubation conditions.

| Antibody | Manufacturer | Clone | Host | Antigen retrieval | Dilution | Incubation |
| --- | --- | --- | --- | --- | --- | --- |
| HER4 | Santa Cruz | C-18 | Rabbit | EDTA pH 8.8 | 1/2500 | Overnight |
| pHER4 (pTyr1162) | Abcam | EP2270Y | Rabbit | EDTA pH 8.8 | 1/100 | Overnight |
| ER | Novocastra | 6F11 | Rabbit | Citrate pH 6.0 | 1/100 | 1h |
| PR | Novocastra | PGR | Rabbit | Citrate pH 6.0 | 1/200 | 1h |
| EGFR | Invitrogen | 31G7 | Rabbit | Chymotrypsin pH7.8 | 1/100 | 1h |
| pEGFR (pTyr1068) | Cell signaling | D7A5 | Rabbit | EDTA pH 8.8 | 1/25 | 1h |
| HER2 | Dako | poly | Rabbit | Citrate pH 6.0 | 1/200 | 1h |
| pHER2 (pTyr1221/1222) | Cell signaling | 6B12 | Rabbit | EDTA pH 8.8 | 1/200 | 1h |
| HER3 | LSBio | 3F10F6 | Rabbit | Citrate pH 6.0 | 1/1000 | 1h |
| pHER3 (pTyr1289) | Cell signaling | 21D3 | Rabbit | EDTA pH 8.8 | 1/50 | 1h |
| YAP | Cell signaling | D8H1X | Rabbit | Citrate pH 6.0 | 1/100 | Overnight |
| pYAP(pSer127) | Cell signaling | poly | Rabbit | Citrate pH 6.0 | 1/250 | Overnight |

### Immunohistochemistry

Immunohistochemistry was performed on 4 μm thick formalin-fixed paraffin-embedded TMA sections using the Mach1 Universal HRP-Polymer Detection Kit (BioCare Medical, CA, USA) according to the manufacturer’s instructions. Briefly, sections were deparaffinized with xylene and hydrated in a series of graded ethanol (95%-70%) to water. Heat-induced antigen retrieval was performed using a decloaking chamber™(BioCare Medical, CA, USA) with either sodium citrate buffer (0.01 M, pH 6.0) for 5 min at 125 °C, or EDTA buffer (0.001 M, pH 8.8) for 30 min at 95 °C. Enzymatic antigen retrieval was performed using α-chymotrypsin (Sigma-Aldrich, MO, USA) for 10 min at 37°C. Sections were rinsed with Tris-buffered saline (TBS) and then treated with 0.3% hydrogen peroxide for 10 min to block endogenous peroxidases. Non-specific antibody staining was blocked with MACH1 Sniper blocking reagent (BioCare Medical, CA, USA). Primary antibody diluted in TBS was applied to the slide for 1 hour at room temperature or overnight at 4 °C in a humidified slide chamber. For rabbit primary antibodies, MACH1 anti-rabbit secondary antibody conjugated to horseradish peroxidase was applied for 30 min at room temperature. Diaminobenzidine chromogen substrate was applied for 1-5 min. Lastly, slides were counterstained with hematoxylin for 30 sec and coverslipped with DPX mountant (Sigma-Aldrich, MO, USA). For analysis, slides were scanned at 40X using the Aperio AT Turbo (Leica).

### Scoring and Analysis

All scoring was performed by three independent observers (PK-dC, ML, JS) using the iPhoto image software (Apple Inc) in a blinded manner. Only invasive cancer cells were scored. Scoring criteria for each marker has been described in figure 1 and Suppl figure 2. Statistical analysis was performed using PRISM Software (v7) (La Jolla, CA, USA). Associations between HER4 and other markers and clinicopathological parameters were evaluated. Briefly, association between the expression levels of HER4 and pHER4 were compared to (i) other HER family members in both cohorts (ii) YAP and pYAP expression in both cohorts and (iii) breast cancer specific subtype expression were also investigated for each marker in both cohorts. We used paired t-tests, Wilcoxon signed-rank test for paired non-parametric variables. ANOVA, Chi-square and Fisher’s exact tests were performed to correlate categorical variables. *P*< 0.05 was considered significant.

**Figure 1.**
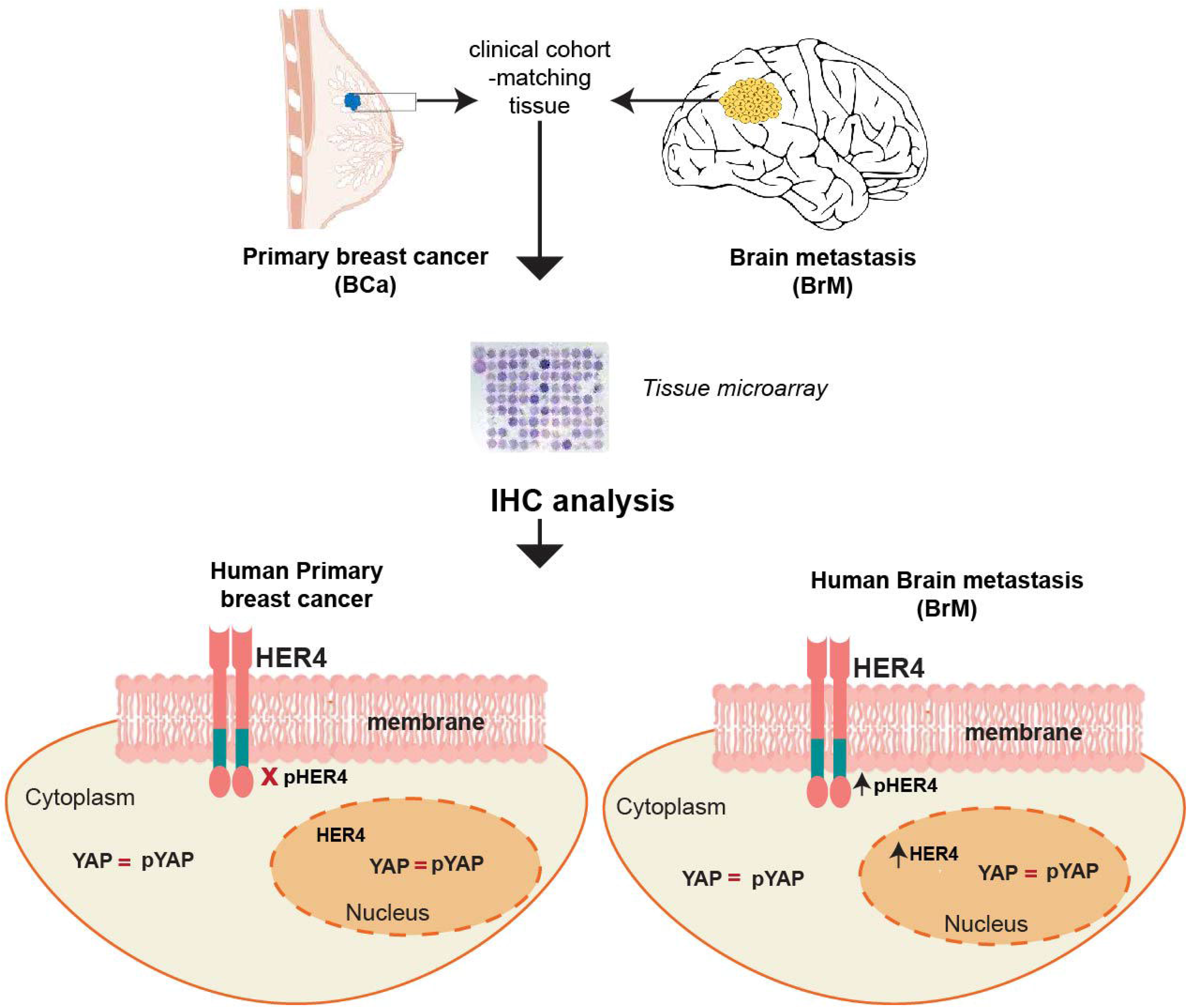
HER4 and pHER4 staining in breast cancers and matched brain metastases. (Ai) Representative images of HER4 expression (top panel) in breast cancer and brain metastasis respectively; pHER4 expression in the membrane is evident in the brain metastatic tissue compared to the breast cancer tissue (bottom right panel). (Aii) Association of HER4 nuclear expression in breast cancer and brain metastasis; χ^2^ (Aiii) IHC staining score of pHER4 in breast cancer compared to brain metastasis; Wilcoxon matched-pairs signed rank test. (B) Heatmap representation of IHC assessment of phosphorylated HER family receptors in breast cancer (BCa) and brain metastasis (BrM). (Ci) Bar graph represents the proportion of cases positive for pHER2 and pHER4 in brain metastasis; stats (Cii) Association of the proportion of cases positive for both pHER3 and pHER4 within brain metastasis; χ^2^. Images were viewed and captured using a 40x objective lens.

## RESULTS

### HER4 expression in brain metastasis is highly activated compared to matched primaries (BBM cohort)

In the breast to brain metastasis (BBM) cohort, the median age at primary diagnosis was 49 years and the median age of brain metastasis diagnosis was 52 years. Median survival after brain metastasis diagnosis was 11 months. HER4 and pHER4 staining was successfully analysed in 37 breast cancer samples and 30 brain metastases samples because of falling out of the cores over time. For YAP and pYAP, 31 breast cancers and 32 brain metastasis cores were scored respectively. The Queensland follow-up cohort has been described in detail by us previously^37^.

Total HER4 protein expression was present in breast cancer (BCa) as well as brain metastasis (BrM) (fig 1Ai) and it was localized in the nuclear and cytoplasmic components of the tumour cells (Suppl fig 1B). Interestingly, BrM samples showed higher HER4 nuclear positivity compared to the primaries (χ^2^, p=0.0043; fig 1Aii). Activation of HER4 was evident with membranous pHER4 staining and nuclear negativity of HER4 suggested HER4 mainly performing its tyrosine kinase activity (fig 1Ai) which was quantified by an IHC score. There was significant increase in the membranous expression of pHER4 in the brain metastasis samples than their primary breast tumours (p=2×10^-6^; fig 1Aiii). Majority of the tumours displaying increased HER4 activation were either HER2+ or TN. Paired t-tests showed significant increase of pHER4 in the matched brain metastatic tissue within the HER2+ and TN group but not in the ER+ group (Suppl fig 1C).

We then investigated the expression of the activated form of the other dimerizing partners^38^ of HER4 including EGFR, HER2 and HER3 in the same patient samples however, a few of them were not matched pairs. For primary BCa, most cases were negative for all HER family partners (fig 1B; top panel). However, in the BrM samples pHER2, pHER3 and pHER4 expression were markedly increased (fig 1B, bottom panel). When the associations between the expression of these proteins were investigated in the BrM group, we found significant increase in the expression of pHER4 with pHER2 and pHER3 (χ^2^, p=0.0057; fig 1Ci and p=0.006; fig 1Cii, respectively).

### HER4 is highly activated in HER2 positive breast cancer (QFU cohort)

Staining of HER4 was both nuclear and cytoplasmic in the general breast cancer cohort (QFU; 95% positivity, 357 out of 373 cases, in all subtypes of breast cancers). Interestingly, pHER4 was positive and membranous in only 16 cases out of 270, with few cases displaying moderate positivity (10) and few showing strong staining (6) (fig 2A). As observed previously in figure 1B, activation of HER4 was significantly associated with its dimerizing partners in brain metastasis. We therefore, investigated whether HER4 activation was enriched in any subtype of breast cancer in the unselected cohort. Although only 6% of the cases (16 out of 270) displayed HER4 activation, 12 cases (75%) were HER2 positive (diagnostic and prognostic criteria included χ^2^, p=2.5 x 10^-8^; fig 2Bi and χ^2^, p=9.3 x 10^-6^; fig 2Bii, respectively). Furthermore, the cases that were strongly pHER4 positive were also highly proliferative as demonstrated by Ki67 staining (antibody already described in^37^; χ^2^, p=0.013; fig 2Biii). Expression of pHER4 expression and total HER4 nuclear positivity remained enriched in brain metastasis compared to all breast cancers including their matched primaries (BCa-BBM) (One-way ANOVA, p=1.2 x 10^-10^; fig 2Ci; p=0.043; fig 2Cii). HER4 cytoplasmic expression was not found to be significantly different between the groups. (fig 2Ciii). Overall, our results indicate that HER4 is highly phosphorylated in HER2+ve breast cancers.

**Figure 2.**
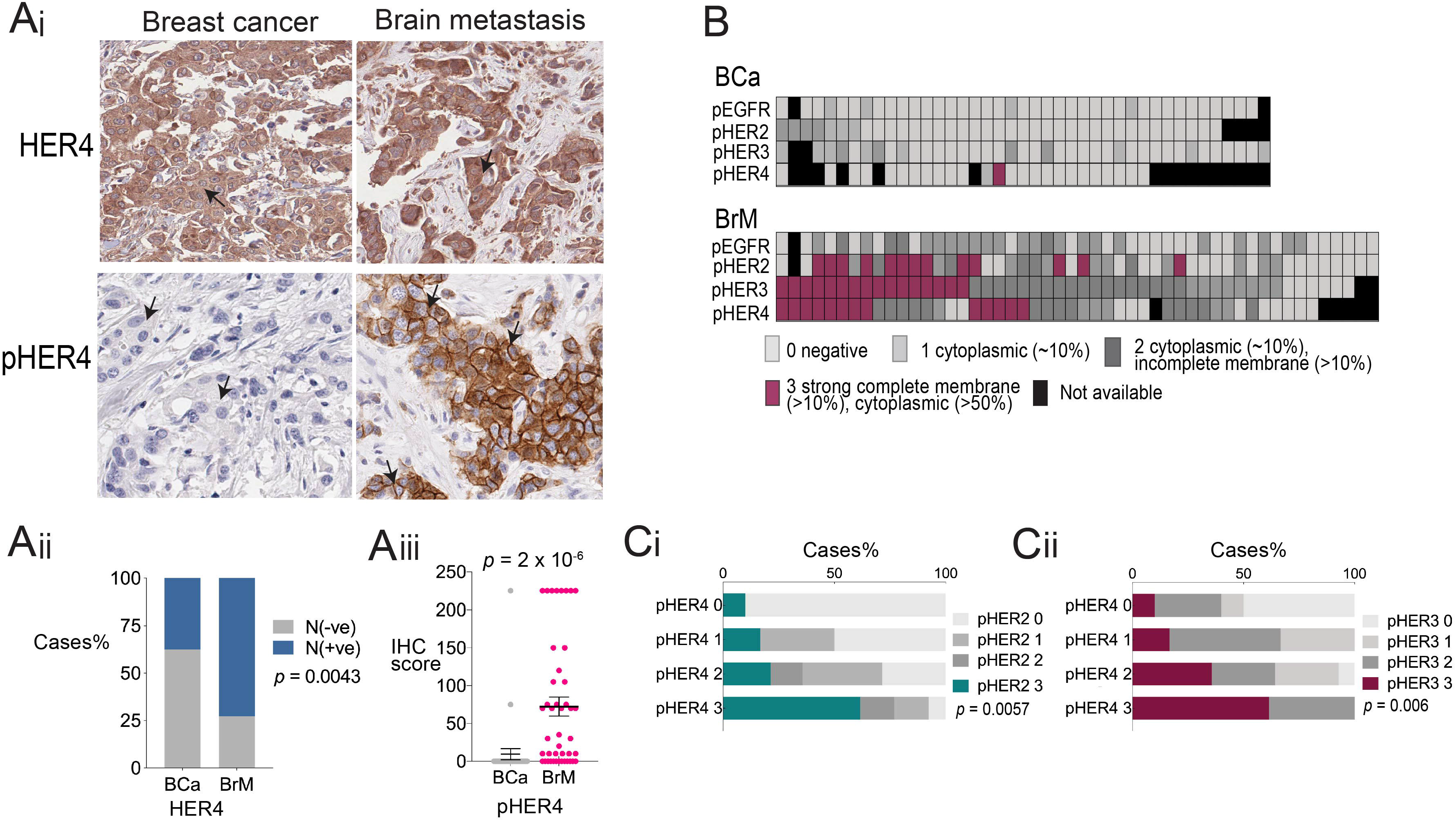
Representative images of the IHC staining of HER4 (Ai) and pHER4 exhibiting moderately (Aii) and strongly (Aiii) positively stained membranes in the QFU cohort (A) coupled with the magnified regions (Bottom panels respectively). (Bi) Association of pHER4 membranous expression in HER2 positive disease (Dx) as well as prognostic criteria (Bii) where again HER2 positivity is enriched with HER4 activation; χ^2^. Proliferative index indicated by Ki67 expression suggests strong association of HER4 activation in HER2+ve disease with highly proliferative tumours (Biii); χ^2^. (C) bar graphs represent the comparison of HER4 and pHER4 expression in breast cancers and brain metastasis within two different cohorts. Cytoplasmic HER4 expression is more widely distributed between all the cases irrespective of the groups and subtypes (Ci); χ^2^. Nuclear expression of HER4 is shown in (Cii) where brain metastasis has relatively higher expression compared to its matched primaries. Remarkable activation of HER4 is seen in brain metastasis compared to all the other groups (Ciii). Images were viewed and captured using a 4x and 40x objective lens.

### YAP protein is significantly phosphorylated in breast cancers and brain metastasis matched pairs (BBM cohort)

The dynamic activity of HER4 after undergoing intramembrane proteolysis, which releases a soluble intracellular domain, has been shown to activate the Yes-associated protein (YAP) in the nucleus (in-vitro)^23, 39^ (fig 2A). The antibody that was used is mapped to the C-terminus of human HER4. Although it does not differentiate between the cleaved and the uncleaved form of HER4, it still recognizes the intracellular domain. This is evident because of nuclear staining of HER4 which is exclusively shown to be the intracellular domain^16–22^. Based on evidence from the literature, we undertook an assessment of the expression of YAP and pYAP (ser127) in our cohorts to determine if there was any association of expression levels of YAP and HER4 proteins in brain metastasis.

We first analysed the expression of YAP and pYAP along with their association with HER4/pHER4 expression in the BBM cohort. We found YAP and pYAP to localize in the cytoplasm as well as the nucleus of the tumours cells with the majority of cases positive for both markers (Supp fig 2A). There was no difference observed in the expression of these proteins in BCa and BrM visually (fig 3Bi) as well as statistically (Suppl fig 2Bi-iv). To test the specificity of the total protein antibodies we validated them using transient knockdowns in breast cancer cells (Suppl fig 2C). The phosphorylation of YAP was strongly associated with the total YAP expression in the cytoplasm (p=0.0366; fig 3Bii) as well as in the nucleus (p=0.0323; fig 3Biii) of BCa and BrM (p=0.0073; fig 3Biv and p=0.0313; fig 3Bv respectively). Within the brain metastasis cases, we investigated the association between HER4 expression and YAP expression but we did not find any direct correlation in the expression of these proteins. However, we did observe HER4 cytoplasmic expression to be significantly associated with YAP cytoplasmic localisation (p=0.0113; fig 3Ci), although those cases were mostly phosphorylated for YAP (p=0.0026; fig 3Cii). Interestingly HER4 cytoplasmic staining also inversely correlated with YAP’s nuclear phosphorylation (p=0.003; fig 3Ciii). Previously, the nuclear expression of HER4 in BrM was found to be higher as per fig 2Aii, therefore we investigated if YAP’s nuclear localisation had any relation to this finding. We found no significant association between HER4 nuclear and YAP nuclear localisation (p=0.1208; fig 3Civ) although 14 cases out of 17 (82%) in the HER4 nuclear positive group had YAP nuclear positivity as well (fig 3Civ). Furthermore, as we saw a strong association between YAP expression and its phosphorylation (fig 3B), we hypothesized that the majority of the cases that are HER4 nuclear positive and YAP nuclear positive might have phosphorylated YAP. Interestingly, as shown in fig 3Cv, comparing HER4 nuclear, YAP nuclear and pYAP nuclear staining, 8 out of 14 cases are phosphorylated. When we analyzed this using chi-square, although it did not reach statistical significance, we found that 6 cases (57%) in the HER4 and YAP nuclear (N) positive category to be phosphorylated compared to only 2 cases (33%) in the HER4 and YAP(N) negative category (p=0.5765; fig 3Cv). We also analyzed the mRNA levels of ERBB4 and YAP1 in our in-house dataset^9^ to detect any differences in the expression between brain metastasis arising from different primary cancers. We found ERBB4 expression to be higher in brain metastasis arising from breast cancers compared to those arising from melanomas (p=0.0313; fig 3Di) whilst they were comparable to those arising from the lung. Interestingly for YAP expression we found lung-BrM to have the highest YAP1 expression compared to melanoma-BrM but comparable to breast-BrM (p=0.0083; fig 3Dii).

**Figure 3.**
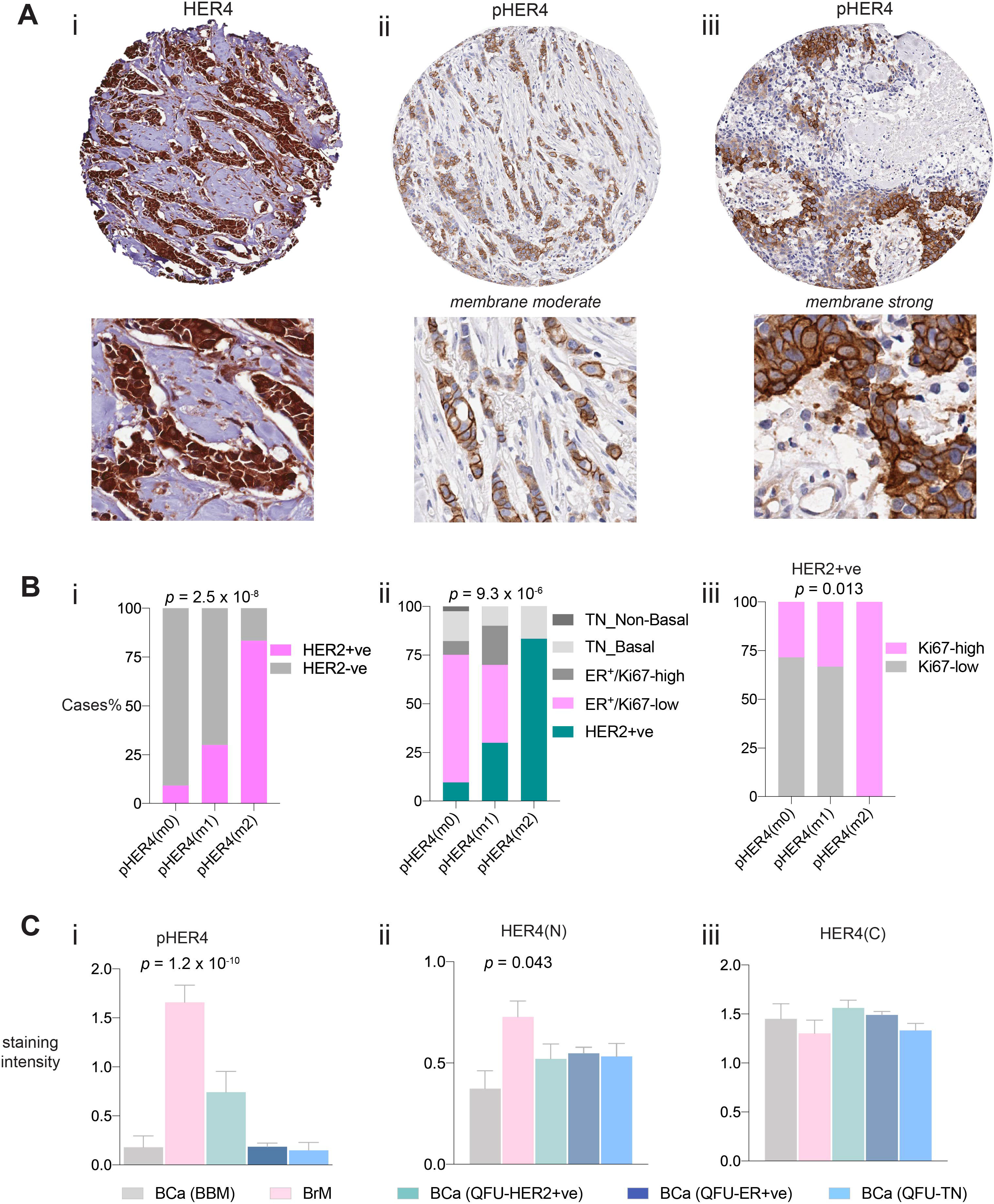
Schematic of YAP and HER4 interaction shown by in-vitro studies based on Haskins et al^23^. Proteolytic cleavage of HER4 resulting in the formation of the intracellular domain and subsequent translocation into the nucleus which results in the activation of YAP, promoting transcriptional activity (A). Immunohistochemical analysis of YAP and pYAP on breast and brain metastasis tissue (Bi). Chi-square analysis of the staining intensities in breast cancer (BCa) between YAP and pYAP in the cytoplasm (Bii) and the nucleus (Biii). Bar graphs of Chi-square analysis of the brain metastasis of the same proteins in the cytoplasm (Biv) and nucleus (Bv). Chi-square association of HER4 cytoplasmic expression with YAP cytoplasmic expression (Ci); pYAP cytoplasmic expression (Cii); pYAP nuclear expression in brain metastasis. Bar graph (Civ) shows association of HER4 nuclear positivity to YAP nuclear positivity using chi-square analysis. (Cv) is a heatmap representation of the brain metastasis cases displaying the positivity of HER4, YAP and pYAP in the nucleus and the bar graph on the right shows association of dual positive or negative cases (HER4/YAP-nuclear) with pYAP nuclear expression. mRNA values for HER4 gene (*ERBB4*) (Di) and YAP gene (*YAP1*) (Dii) are displayed in the scatter dot plot graphs in brain metastasis cases arising from three different primary cancers. Images were viewed and captured using a 40x objective lens.

### YAP protein is highly phosphorylated in breast cancers (QFU cohort)

We further investigated the expression of YAP and pYAP in primary breast cancer cases from the unselected cohort (QFU). Both proteins showed positive expression in the nucleus and the cytoplasmic in all breast cancers (fig 4A). We examined the sub-cellular localisation of YAP and pYAP in ER+ve, HER2+ve and triple-negative breast cancers. Surprisingly, we found that irrespective of the cellular compartment, YAP is highly phosphorylated in ER+ve (p=1 x 10^-15^; fig 4Bi and p=2 x 10^-13^; fig 4Bii) and HER2+ve breast cancer (p=0.0009; fig 4Biii and p=8.5 x 10^-4^; fig 4Biv) as well as TNBC (p=2.7 x 10^-10^; fig 4Bv and p=2.9 x 10^-5^; fig 4Bvi). Interestingly we found no association between HER4 expression and YAP expression in HER2+ve and TNBC, however, within ER+ve disease, we discovered that cases with high HER4+ve cytoplasmic staining were also significantly positive for YAP cytoplasmic expression (p=0.022; fig 4Ci). However, these cases were also significantly associated with YAP’s phosphorylation (p=5 x 10^-13^; fig 4Cii). Similarly, the nuclear compartment showed complementary results with HER4 nuclear positivity associating with YAP’s strong nuclear expression (69 cases/79%, p=3 x 10^-10^; fig 4Ciii). When we investigated the association between nuclear YAP expression with pYAP expression within the HER4 nuclear positive cases, 43 cases (68%) also displayed strong pYAP positivity (p=4.6 x 10^-5^; fig 4Civ). Utilizing the data generated as part of previous studies^37, 40–42^, we further investigated the association of YAP’s nuclear positivity with its known downstream effector, the transcription factor SOX9^43^. We found a significant association between YAP nuclear positivity and the nuclear expression of SOX9 within TNBC (p=1.5 x 10^-5^; fig 4Di), both basal-like (p=0.0036; fig 5Dii) and non-basal (p=0.0476; fig 4Dii). Our results also indicated that this association is perhaps functionally active as 12 cases (42%) of the YAP nuclear positive cases were not phosphorylated within the SOX9 nuclear positive tumours (p=0.07; fig 4Div).

**Figure 4.**
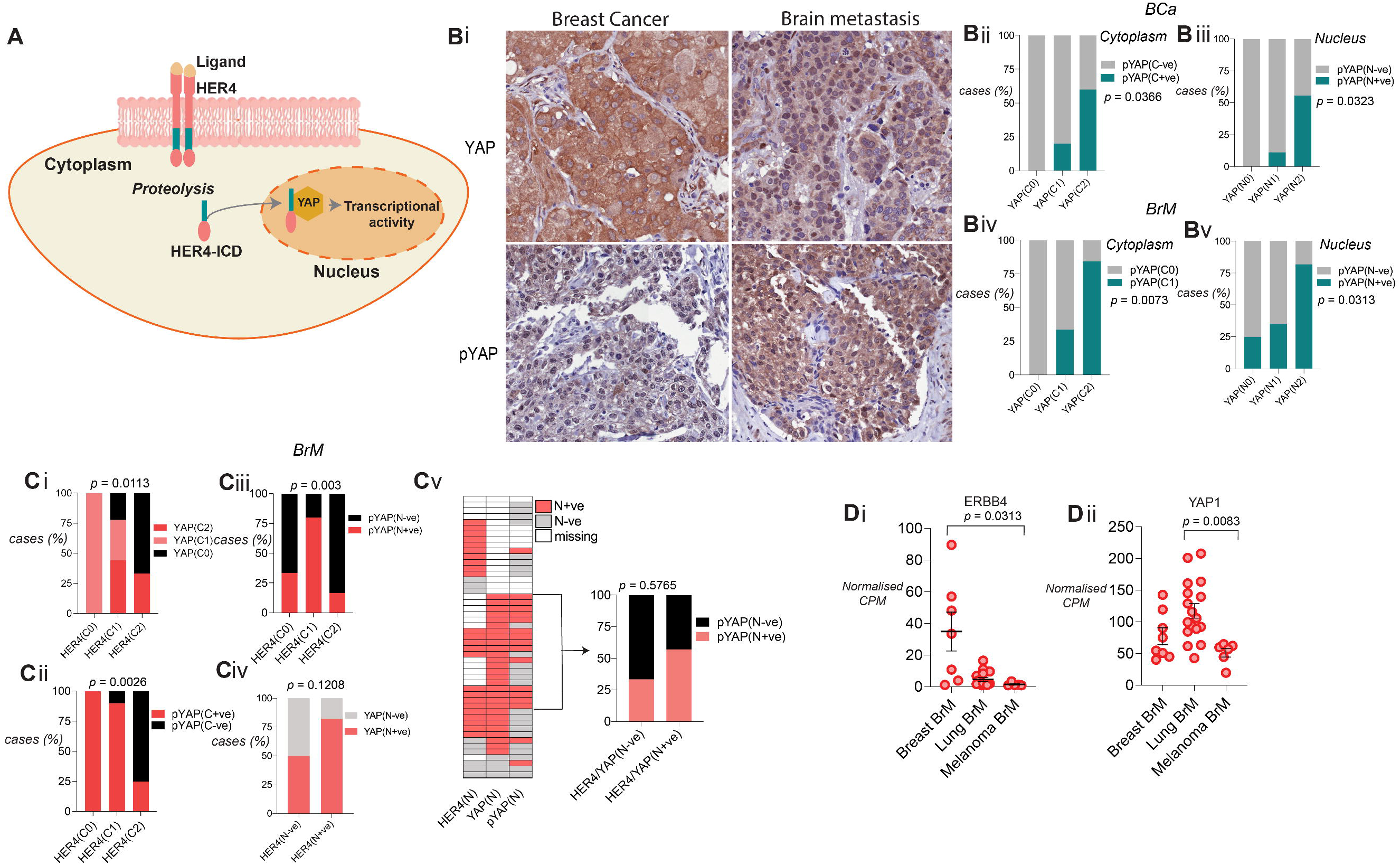
Positive staining of YAP and pYAP on primary breast cancers (A). (B) Chi-square analysis of the staining intensities in the three subtypes of breast cancer between YAP and pYAP in the cytoplasm (i, iii, v) and the nucleus (i, iv, and vi). (C) Bar graphs of Chi-square analysis within ER+ve tumours show strong association between HER4 cytoplasmic positivity with YAP cytoplasmic localization (i) which are also highly phosphorylated (ii). A similar trend was observed in the nuclear staining of these tumours with HER4 significantly associating with YAP (Ciii) and these tumours were highly phosphorylated for YAP (C iv). A different association with respect to YAP was observed within TNBC where the transcription factor SOX9 associates significantly with YAP nuclear localization (Di) irrespective of the tumours being basal-like (D ii) or non-basal (D iii). (D iv) this bar graph illustrates that 42% of the tumours within the SOX9 positive group are not phosphorylated for YAP. Images were viewed and captured using a 4x and 40x objective lens.

## Discussion

The present study demonstrated a significant upregulation of phosphorylated HER4 in matched-pair, clinical brain metastasis samples. This activation was significantly associated with HER2 and HER3 activation in brain metastasis. Furthermore, in primary breast cancer, activation of HER4 strongly associated with HER2 positive subtype. These results suggest that there might be a positive feedback loop where activation of HER2 feeds into the other partners of the pathway. This is plausible given that some of the ligands are common between the different dimer partners. With respect to HER4/YAP axis, surprisingly we found no association of YAP’s expression with HER4 expression and activity suggesting HER4/YAP axis might not be active in breast cancer-BrM. In addition, irrespective of the primary or the metastases YAP was highly phosphorylated suggesting that YAP might be in an inactive state in the majority of the tumours. Interestingly, in TNBC, YAP’s active form in the nucleus strongly associated with the transcription factor SOX9 expression suggesting YAP-driven transcriptional activity and these findings concurred with the results from another study as previously reported^43^.

The brain specific adaptations that set BrMs apart from their primary cancers are evident from various studies^44, 45, 46, 47–49^, where the tumour cells engineer peculiar ways to adapt to this hostile environment. From creating communicating bridges between neuronal cells and themselves^44^; mimicking the functions of neurons^48^; releasing factors that would curtail the defensive program of the neurons^45^; to using unique metabolic proteins expressed in the TME to aid their growth^49^; tumour cells are skillful warriors in establishing their colonies. The findings from this study revealed similar results where HER4 activation is more significant in the brain metastases compared to their parent breast tumours. Neuronal cells express HER4 abundantly for various functions^50^, in addition one of its ligand NRG-1 is also present in vast amounts within the brain^11^. We hypothesize that the tumour cells which express significantly high phosphorylated HER family receptors are merely taking advantage of this ligand-rich microenvironment; which ultimately facilitates the growth of the tumour. To the best of our knowledge, this is the first study to report this change in a matched-pair series.

HER4 has been shown to mediate trastuzumab resistance in HER2 +ve BC^31^ and gastric cancer^51^. In HER2+ve gastric cancer, it is shown to drive the resistance to trastuzumab via mediating epithelial to mesenchymal transition through its downstream effector YAP1 protein. We did not find any correlation between YAP and HER4 in HER2 +ve disease in our cohorts. This is not a surprising finding as our patient groups were treatment naïve, meaning these tumours were resected pretreatment. Furthermore, the general breast cancer cohort is a historical cohort meaning these patients were diagnosed with breast cancer before trastuzumab was approved as a standard first-line therapy for HER2+ve breast cancer^52, 53^ and neoadjuvant therapy was not common in that era. This reflects how the treatment course of a disease can impact these pathways and future studies should investigate tissues from treatment resistant or incomplete/partial pathological response groups from neoadjuvant therapy cohorts for the HER2+ve patients. Nonetheless, our results clearly suggest that HER4 activation in primary breast cancer is more common in HER2+ve disease. Therefore, combination therapy targeting multiple HER receptors might be a feasible option. Recent Phase I and II clinical trial studies indicate that a new generation pan-HER inhibitor Pyrotinib (EGFR, HER2 and HER4) might be a potential drug for HER2+ve metastatic breast cancers. The Phase I study reported acceptable efficacy and tolerability of this drug in metastatic HER2+ve breast cancers^54^. This was followed by the Phase II trial, where pyrotinib was prescribed and tested against lapatinib (EGFR and HER2 inhibitor) with a combination of capecitabine (antimetabolite chemotherapeutic agent)^55^. The Pyrotinib arm displayed a better response (~79%) compared to ~57% in the lapatinib arm and the median progression free survival also was longer for pyrotinib treated patients, thus, providing feasibility for the following Phase III trial. Therefore, this study provides a rationale for a combination treatment with small-molecule inhibitors together with chemotherapeutic agents for HER2+ve metastatic breast cancers.

YAP is a key regulator of the hippo tumour suppressor pathway and is being researched quite extensively in breast cancer. The contextual dependency of YAP’s function as either a tumour promoter^28, 56–60^ or a tumour suppressor^27, 61–64^ in cancer is widely reported. With respect to breast cancer, one study reported a decrease in YAP levels and genomic depletion of YAP from breast cancer cell lines suppressed anoikis and increased the migratory properties of the cells^63^. Similarly, Cao and colleagues showed that at the protein level expression of YAP in 324 breast cancer patients had a favorable prognosis especially in the PR subgroup and Luminal A prognostic subtype. In addition, YAP expression and PR status were found to be independent predictors of disease-free survival and overall survival^27^. It should be noted that this cohort is a contemporary cohort ranging from between 2009 to 2014 compared to our unselected cohort, so current therapies might have also influenced the outcome of these patients. In contrast, the expression levels at the mRNA of *YAP* in breast cancer have been recently investigated and it was found to be a poor prognostic factor. Using in-vitro techniques the study showed that YAP promotes cell growth and decreases apoptotic capability of breast cancer cells^28^. As far as we know, our findings are completely unreported in the literature; with YAP being highly inactive in breast cancer and brain metastasis. This is a novel and confounding finding as previous studies including in-vitro and clinical samples have shown contradictory results^23, 25, 27, 28, 65^. This leads us to speculate that perhaps there is an unknown function of phosphorylated YAP yet to be uncovered and its regulation in the nucleus is poorly understood. Recently, the mRNA splicing nuclear kinase PRP4K was shown to phosphorylate YAP in the nucleus^66^. Kinome screen mediated analysis revealed PRP4K to increase YAP expression which acted in parallel with the other hippo pathway proteins. This not only restricted its binding affinity but also excluded YAP from the nucleus. This mechanistic insight was reported in Drosophila, however, the authors also conducted in-vitro studies to elucidate the implications of this finding in TNBC. They found that the transient knockdown of PRP4K decreased YAPs phosphorylation and increased total YAP levels in breast cancer cells. This clearly indicates a novel role of this kinase in regulating YAP in breast cancer. However, the authors did not show cellular compartment specific expression of these proteins in breast cancer. Nonetheless, based on this study we can speculate that perhaps in our cohorts this kinase is present and this rigorously phosphorylates YAP in the nucleus and expels YAP back into the cytoplasm and therefore, we observe equal amounts of YAP along with its phosphorylated form. Future studies should investigate PRPK4 protein in detail in clinical cohorts of breast cancer and brain metastasis cohorts.

HER4 can bind to YAP in its WW domain similar to the inhibitory molecule LATS1. HER4 may reduce inhibitory phosphorylation of YAP by LATS1 by competing to bind. We observed in the majority of the cases with BrM displaying high HER4 positivity along with high pYAP in the cytoplasm. It is plausible to assume that whenever HER4 was expressed highly in the cytoplasm it was mostly activated, meaning it could not bind to YAP hence leaving it available for phosphorylation/ inactivation by LATS1.

Although YAP is mainly inactive in ER+ve breast cancer, we did observe a positive association between HER4 and total YAP in the nucleus. A novel non-canonical functional relationship between YAP and ERα has been recently demonstrated which drives breast cancer growth of ER+ve breast cancer in-vivo and in-vitro^67^. Furthermore, HER4 ICD has been demonstrated to be a co-activator of ERαand it drives its expression and breast tumour proliferation in response to estrogen^17, 68^. Taking together the data from different studies, it is tempting to speculate that perhaps the cases which have high HER4 nuclear positivity along with YAP might be more proliferative. Unfortunately, we did not see any significant difference in Ki67 expression due to small numbers within that group (data not shown).

The association of YAP with SOX9 has broadly been described in gastric^69^ and esophageal cancers^43, 70^. In gastric cancer cell lines, a causal relationship between SOX9 and YAP has been shown; where knocking down SOX9 leads to reduced expression of YAP protein and increases its phosphorylation^69^. The stem-like properties in esophageal cancers are imparted by the YAP-SOX9 axis. This axis has been shown to help the cancer cells acquire cancer stem-cell like properties in-vitro^70^. In another study, inhibition of YAP led to a reduction in cancer stem-cell like properties of esophageal cancer cell-lines and the authors also reported transcriptional activation of SOX9 by YAP. In two independent clinical cohorts, YAP and SOX9 were shown to be significantly associated. This finding is similar to our finding where TNBC had a strong association of YAP and SOX9 expression in the nucleus. SOX9 mRNA expression has been implicated in estrogen-negative breast cancers^71^. In-vitro analysis on TNBC cell lines have described SOX9 to be a driver for invasion and metastasis^72^. Although we did not observe poor prognosis in the patients with co-expression of YAP and SOX9, this association might have endowed stem-like features to those tumours; furthermore, we cannot rule out that if we surveyed a bigger TNBC cohort we would not find prognostic implications of this association.

Overall, HER4/YAP axis expression is not significantly associated in breast cancer brain metastasis suggesting combination therapies using HER4 and YAP inhibitors might not be a feasible option for these cancers. This study highlights the role of HER4 protein in breast cancer brain metastasis especially in HER2 positive disease suggesting a combination therapy for example, a Pan-HER inhibitor may be more efficacious in reducing the tumour burden in advanced disease.

## Author Contributions

Conceptualization, PK-dC and JMS; methodology/experiments-PK-dC, ML, XdL, BWD; writing—original draft preparation, PK-dC; writing—review and editing-all authors; supervision, JMS and SRL; project administration, JMS; funding acquisition, JMS and SRL.

## Acknowledgments

We are thankful to Dr. Amy McCart Reed for helpful discussions on the manuscript findings and Dr. Dominique Ezra for editing the manuscript. We are also grateful to the Brisbane Breast Bank (BBB) especially Mr. Kaltin Ferguson for his assistance in collecting cohort demographics information from BBB and to patients past and present who donate tissue and clinical information for research.

---

**Suppl 1** Representative examples of (A) hematoxylin and eosin stained tumour cores for primary breast cancer (left) and metastatic brain tissue (right) from the same patient. **#** depicts the tumour. Total HER4 protein staining in the nucleus as well as the cytoplasm in primary as well as brain metastasis tissue (green arrows indicate cells with nuclear positivity along with cytoplasmic staining). (C) pHER4 analysis showing staining intensity of matched pairs (breast cancer and brain metastasis) of the three subtypes of cancers, HER2+, ER+ and TNBC. Images were captured using a 4x and 40x objective lens.

**Suppl 2** (A) Heatmap representation of YAP and pYAP nuclear and cytoplasmic staining in breast cancer and matched brain metastasis tissue. Bar graphs show chi-square analysis showing the fraction of cases positive for YAP cytoplasm (Bi), nucleus (Bii), pYAP cytoplasm (Biii) and nucleus in breast cancer and their matched brain metastases. Validation of antibodies for the total proteins HER4 and YAP. Transient transfections with siRNA negative control and siRNA HER4 in T47D breast cancer cell line (Top panel) and siRNA YAP (bottom panel) in MDA-MB-468 breast cancer cell line. Images were captured using a 20x objective lens.

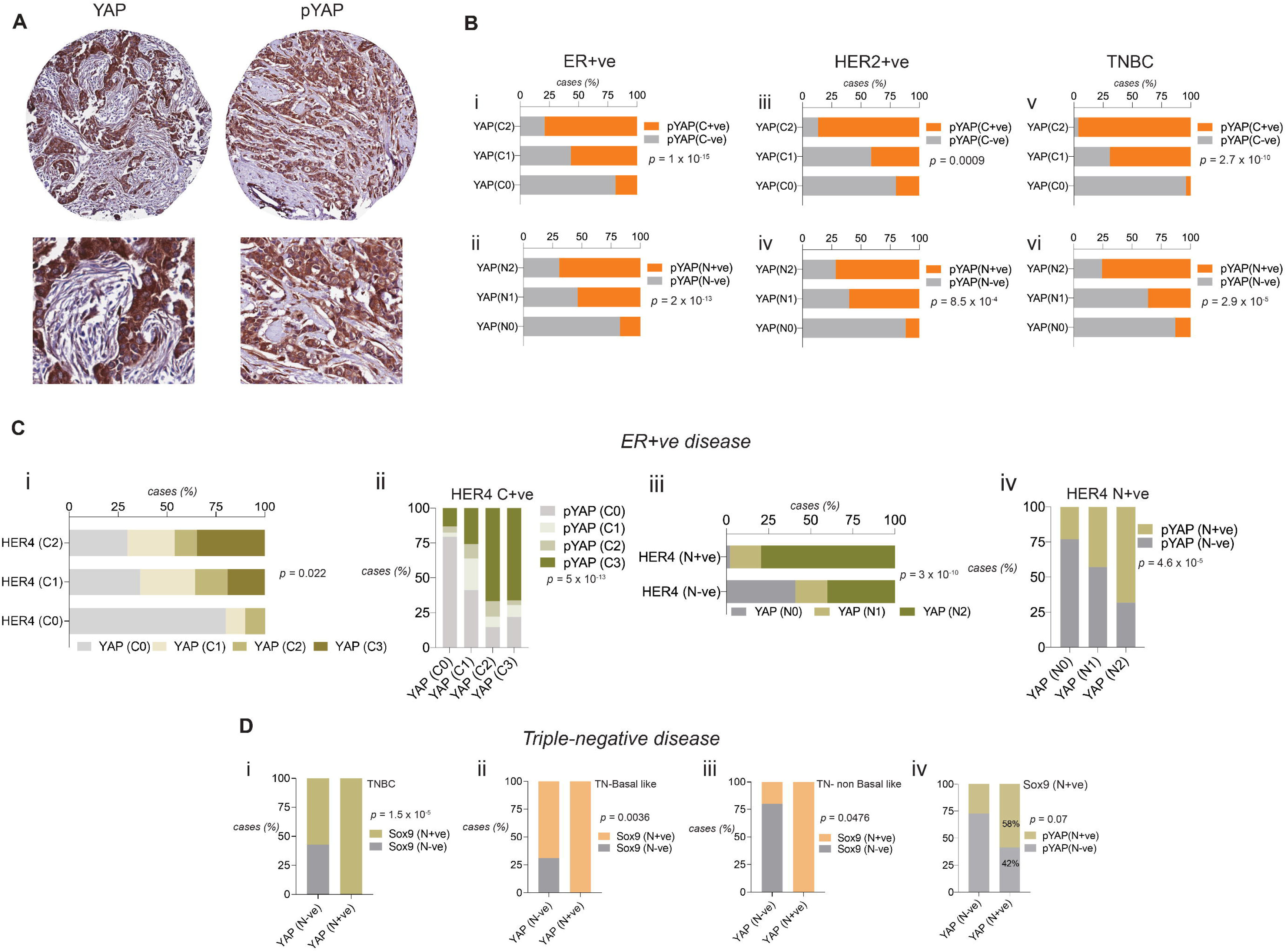

