## Supplementary figures and images for "Immunophenotyping of HER4/YAP axis in breast cancer brain metastasis"

### Supplementary Figure 1

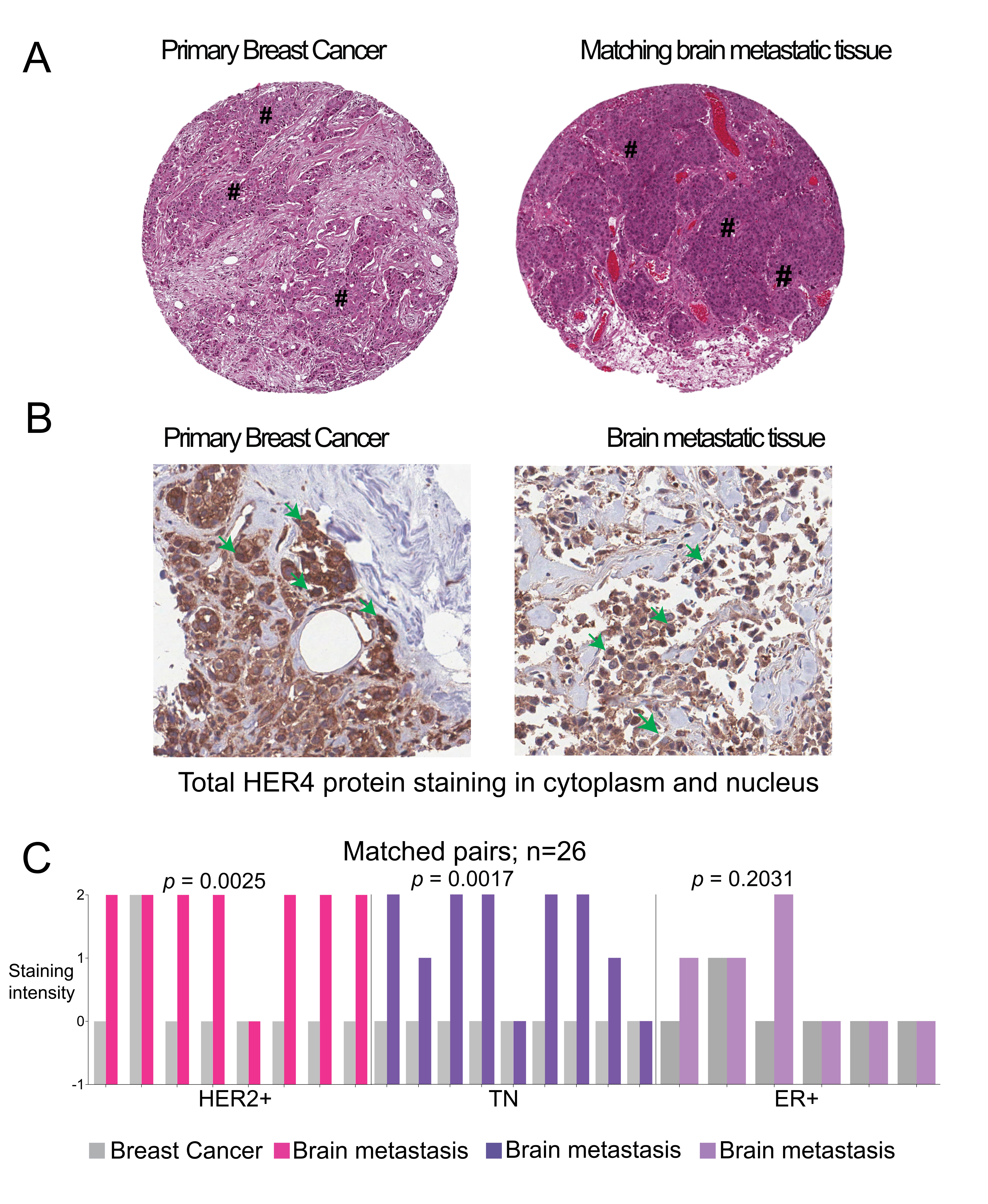

### Supplementary Figure 2

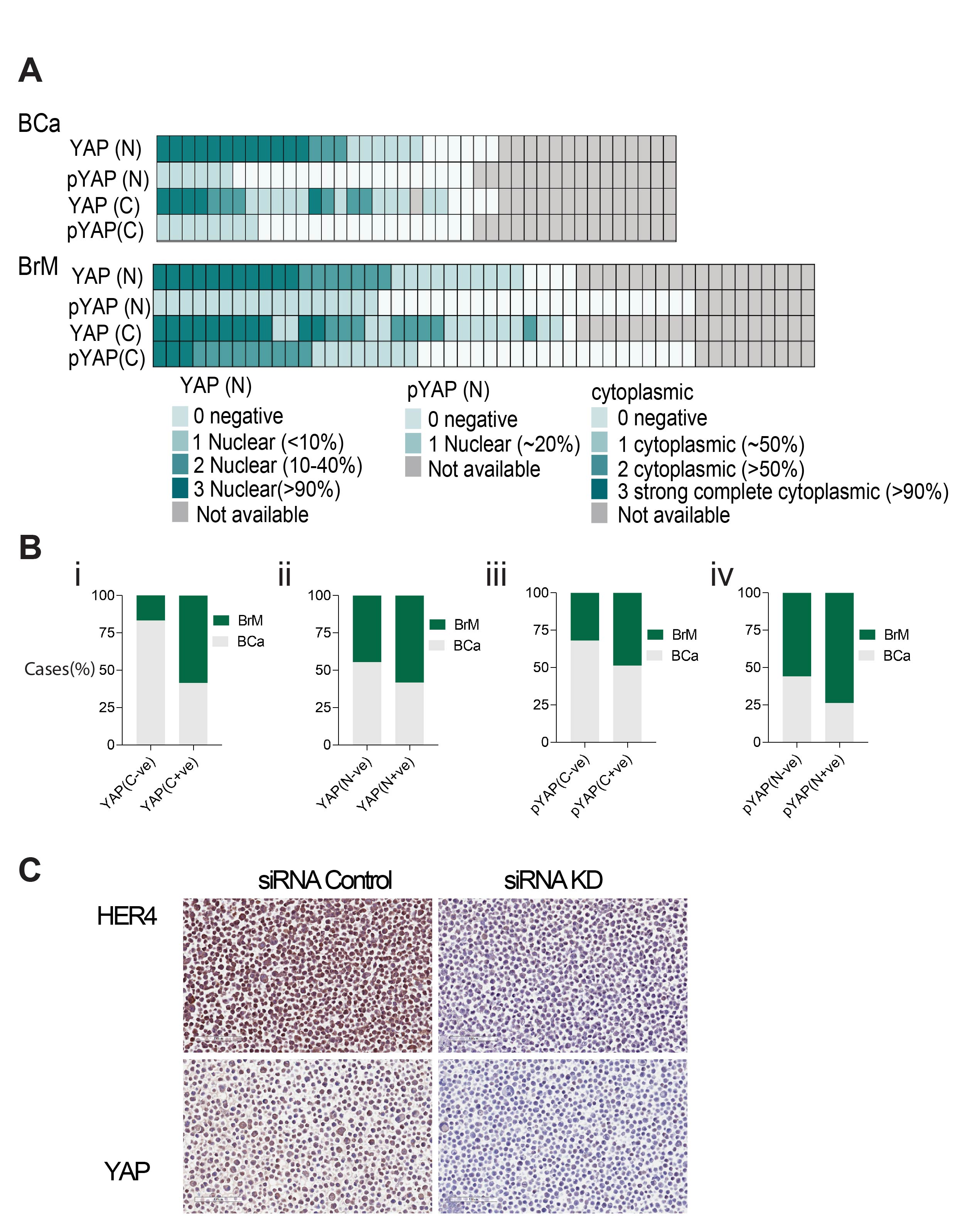
